# Increasing the Field-of-View in Oblique Plane Microscopy via optical tiling

**DOI:** 10.1101/2022.06.12.495844

**Authors:** Bingying Chen, Bo-Jui Chang, Felix Zhou, Stephan Daetwyler, Etai Sapoznik, Gabriel M. Gihana, Lizbeth Perez Castro, Maralice Conacci Sorrell, Kevin M. Dean, Alfred Millett-Sikking, Andrew G. York, Reto Fiolka

**Affiliations:** Lyda Hill Department of Bioinformatics, UT Southwestern Medical Center, Dallas, Texas, USA; Department of Cell Biology, UT Southwestern Medical Center, Dallas, Texas, USA; Calico Life Sciences LLC, South San Francisco, CA, USA

## Abstract

Fast volumetric imaging of large fluorescent samples with high-resolution is required for many biological applications. Oblique plane microscopy (OPM) provides high spatiotemporal resolution, but the field of view is typically limited by its optical train and the pixel number of the camera. Mechanically scanning the sample or decreasing the overall magnification of the imaging system can partially address this challenge, albeit by reducing the volumetric imaging speed or spatial sampling, respectively. In this Letter, we introduce a novel dual-axis scan unit for OPM that enables rapid and high-resolution volumetric imaging throughout a volume of 800 × 500 × 200 microns. This enables imaging of model organisms, such as zebrafish embryos, with subcellular resolution. Furthermore, we combined this microscope with a real-time and multi-perspective projection imaging technique to increase the volumetric interrogation rate to more than 10 Hz.

---

Recent developments in Oblique Plane Microscopy (OPM) [1-3] have demonstrated its potential for 5D (XYZλT) live imaging of cells and small model organisms. Like Light-Sheet Fluorescence Microscopy (LSFM), OPM leverages a highly parallelized acquisition and detection scheme, which greatly reduces photobleaching [4]. Moreover, it uses a single objective for both illumination and detection, which alleviates the imaging space constraint inherent to most LSFMs, reduces the complexity of sample preparation, and brings great potential in high throughput volumetric imaging [5]. Lastly, rapid volumetric imaging can be achieved in OPM systems using rotating polygon mirrors [3] or galvanometric scanners [6, 7] that scan the light-sheet and de-scan the fluorescence light.

However, OPM systems to date have a limited field-of-view (FOV), mainly due to their complex optical detection train. To address this challenge, the FOV has been increased by mechanically scanning the stage [8] and/or decreasing the overall magnification of the imaging system [8, 9]. However, these approaches lead to either reduced volumetric imaging speed or a compromised sub-Nyquist spatial sampling rate. Here we introduce an optical tiling OPM system that enables rapid volumetric imaging of large live samples while maintaining Nyquist sampling.

The FOV limitations in OPM are rooted in its remote focusing mechanism, which is necessary to create a distortion free image of the tilted light-sheet plane [10]. To work properly, the intermediate image in between the secondary and tertiary objective [termed O2 and O3 in Fig. 1(a)] needs to be magnified by the ratio of the refractive indices between the sample space and the remote image space [10]. When air objectives are used for the secondary objective, necessary to enable recent improvements in resolution and light collection in OPM [2, 11, 12], typically a magnification of 1.33-1.51 is required, depending on the choice of the primary objective O1. Thus, the FOV supported by the tertiary imaging system gets demagnified by the same factor. Furthermore, due to the fixed magnification requirement between O1 and O2, the tertiary imaging system must perform the bulk of the magnification to satisfy the Nyquist criterion for spatial sampling. For these reasons, the tertiary imaging system forms a bottleneck for imaging large FOVs, whereas in principle low magnification objectives supporting large FOVs can be used as O1 and O2.

**Fig. 1.**
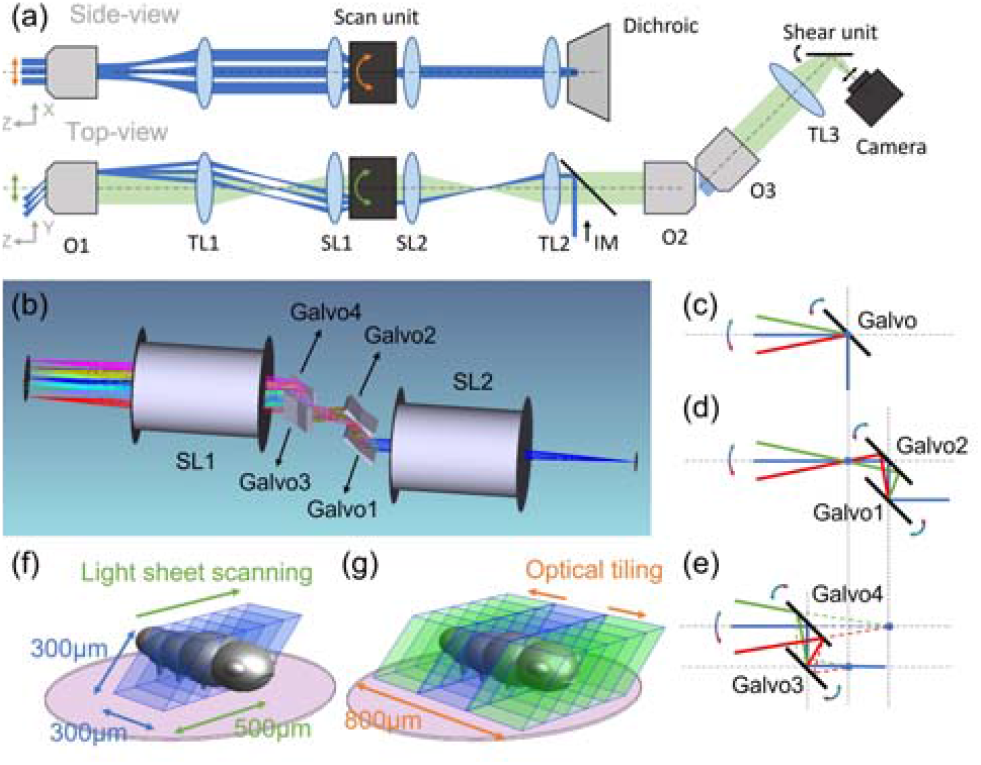
(a) Schematic drawing of the OPM system with optical tiling. O1-3, objectives; TL1-3, Tube Lenses; SL1-2, Scan Lenes, IM, Illumination Module. (b) Rendering of the optical layout of the dual-axis scan unit and scan lenses. (c)-(e) Working principle of a single-galvo, dual-galvo and quad-galvo scan unit. (f) Image volume covered by conventional OPM (blue volume). (g) Image volume covered by optical tiling OPM (blue and green volumes). The pink circle depicts the field of view of the primary objective.

In this study, we aim to utilize the large FOV provided by a 25X NA1.1 water-dipping O1 (Nikon, MRD77220) via a novel, two-dimensional light-sheet scanning approach. To enable telecentric 2D scanning in sample space, typically two galvo mirrors need to be located conjugate to the pupil plane of the primary objective. If one were to insert a conventional, orthogonal dual-axis galvo pair in the space between the two scan lenses (Fig. 1a), only one galvo, or neither, will be conjugated properly to the pupil plane of the primary objective. Importantly, non-telecentric scanning in OPM will alter the tilt angle of the light-sheet in that direction, thereby complicating interpretation of the ensuing data. One way to solve this problem is to add a relay lens pair between the orthogonal galvos. However, this will increase the complexity, and reduce the light-efficiency of the whole imaging system.

To address this challenge, we designed a compact optical dual-axis scan unit, which is based on the original idea that two mirrors can steer a beam through an arbitrary point in space at different angles. Specifically, our scan unit consists of two galvo pairs with 15 mm distance between each galvo (GVS211, Thorlabs), as shown in Fig. 1(b). Pair 1, which includes Galvo 1 and 2, is placed orthogonal to Pair 2 (Galvo 3 and Galvo4). When the scanning angle of Galvo 2 is twice the scanning angle of Galvo 1, the laser beam will pass through a common point [Fig. 1(d)] and the distance between the point and Galvo 2 will be the same as the distance between the two galvos. When this spot lies in the focal plane of the scan lens, the resulting scanning is analogous to a single galvo conjugated to the pupil plane of the primary objective [Fig. 1(c)]. For Pair 2, when the scanning angle of Galvo 3 is set to twice the scanning angle of Galvo 4, a virtual scan point [Fig. 1(e)] is formed that also acts in the same way as a properly conjugated single galvo mirror. By placing the common point in Pair 1 and the virtual point at the focal plane of a scan lens, we can realize dual-axis telecentric scanning with less lenses. A detailed analysis can be found in Section 1 of Supplement 1.

A schematic of our OPM with the dual-axis scan unit is shown in Fig. 1(a). The optical system consists of three major components: the remote focusing system, the illumination module (IM) and the tertiary imaging system. The remote focusing system contains primary objective O1, tube lens TL1 (TTL200, Thorlabs), tube lens TL2 (169 mm Lens assembly [12], TTL200MP and AC508-750-A, Thorlabs) and the secondary objective O2 (20X, air, Olympus, UPLXAPO20X). Two scan lenses SL1, SL2 (CLS-SL, Thorlabs) and the dual-axis scan unit [see also Fig. 1(b)] are inserted between TL1 and TL2 to scan the light-sheet and de-scan the fluorescence light. The excitation laser [blue in Fig. 1(a)] from the illumination module (IM) was introduced into the system by a dichroic mirror (Di03-R405/488/561/635-t1-25×36, Semrock) between TL2 and O2. IM consists of a fiber coupled laser module, a Powell lens to generate uniformly illuminated line and an electrically tunable lens to adjust focal position and to optionally enable axially scanned light-sheet microscopy (ASLM [13], detailed in the Section 3 and 4 of Supplement 1. This mode was not used in the data shown in the main manuscript). The remote focus image formed by the fluorescence light [green in Fig. 1(a)] after O2 is collected by the tertiary imaging system, which is tilted by the same angle as the light-sheet in the sample plane. The tertiary imaging system consists of a solid immersion objective O3 (45° tilted, AMS-AGY v2 [8]) and then focused by TL3 (AC508-300-A-ML) onto a scientific camera (ORCA-Flash, C13440-20CU, Hamamatsu). The overall magnification of the imaging system is 43.4, corresponding to a pixel size of ∼150 nm. To enable optical de-skewing and multi-perspective projection imaging, we also placed a galvanometer-based shear unit in front of the camera [14].

Owing to the dual-axis optical scan unit, the sample can be scanned in one direction to obtain a 3D stack [see also “top view” in Fig. 1(a) and Fig. 1(f)] as in conventional OPM. The other scan dimension is used to extend the field of view by optical tiling [see also “side view” in Fig. 1(a) and Fig. 1(g)], here by acquiring three stacks with different lateral displacements. With this approach we achieve a scan range of 500 µm for a single 3D Stack and extend the lateral width ∼2.5 fold from 300 µm with no tiling [ Fig.1(f)] to 800 µm with optical tiling [ Fig 1(g)]. Importantly, the pixel size of 150 nm assures Nyquist oversampling for an optical resolution around 400 nm. The maximum height of the imaging volume (z-direction) is 200 µm, which is limited both by the length of the light-sheet and the size of the camera.

To characterize the resolving power of our OPM system, we measured the full width half maximum (FWHM) of 100 nm diameter fluorescent nanospheres and performed image decorrelation analysis on subcellular features (Fig. 2). The average resolution across the extended FOV was 0.39±0.05, 0.43±0.04 and 1.22±0.13 µm (n=943) in the x, y and z direction, respectively, as measured by the FWHM [see also Fig. 2(a)]. While the y and z resolution remained relatively constant, the x resolution deteriorated at the periphery of the FOV. This was expected due to some light clipping by the galvo scan unit (see also Fig. S3 in Section 1 of Supplement 1.) Importantly, the z-resolution is better than in a recently published OPM system designed for mechanical tiling with similar optical specifications [8]. This is mainly driven by the detection PSF (i.e. using widefield illumination and OPM detection), which featured an axial FWHM of 1.10±0.15 µm (n=244). We attributed this to the used NA 1.1 primary objective lens, which provides sub-micron axial resolution in a conventional widefield format (not OPM) for green emission. As such we achieve an axial resolution of around one micron while using large light-sheets, required to image multi-cellular specimens.

**Fig. 2.**
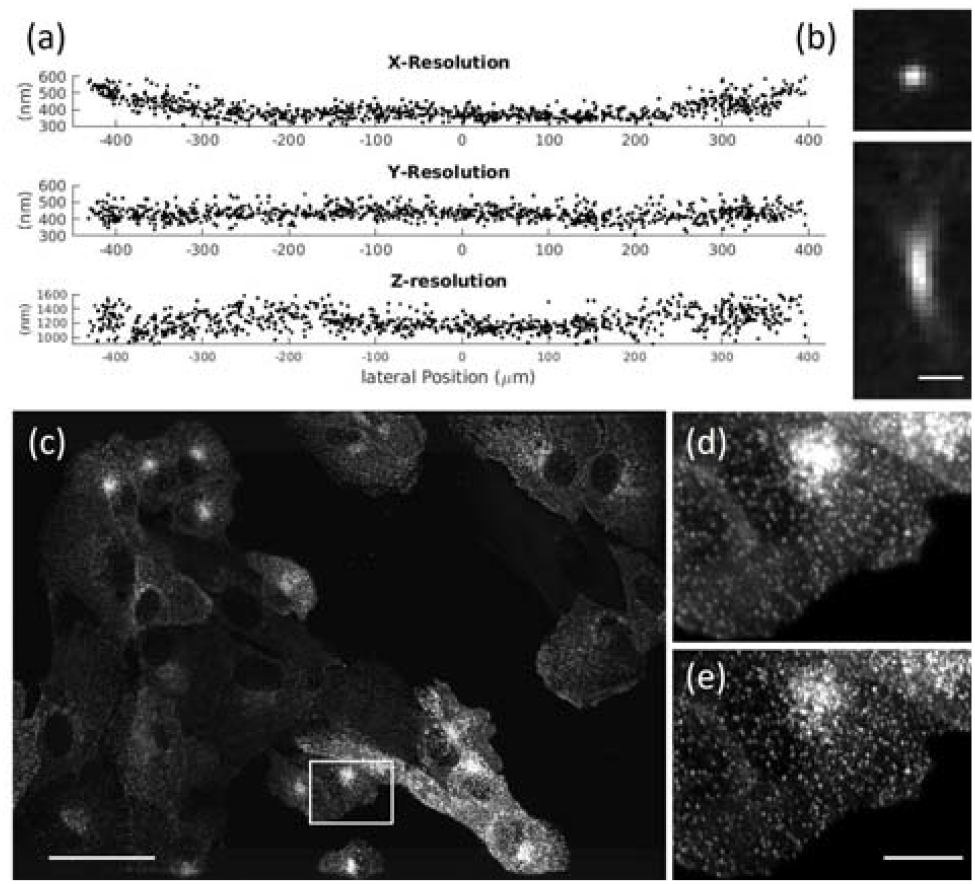
(a) Full Width Half Maximum measurements of fluorescent nanospheres over the whole FOV. (b) Representative point-spread functions of a fluorescent nanosphere. (c) Maximum intensity projection of parental human retinal pigmented epithelium (ARPE-19) cells EGFP-labeled AP2. (d) Raw data for box in (c). (e) Deconvolved data for box in (c). Scale bars: (b)1 µm; (c) 50 µm; (e) 10 µm.

To evaluate the resolving power in a cellular context, we imaged clathrin coated pits. Fig. 2(c) shows a group of ARPE cells labeled with EGFP for AP2 [15], as imaged with our OPM system, while the insets in Fig. 2 (d-e) show magnified regions before and after deconvolution. Using image decorrelation analysis on the whole image, we could verify that in a cellular context we achieve 408 nm resolution in the raw data, and 294 nm resolution after deconvolution (Data was resampled onto a finer grid prior to deconvolution. See also Fig. S10 in Section 5 of Supplement 1).

To demonstrate the spatiotemporal resolution of the system, we next segmented and meshed the 3D volumes of single cells and multicellular constructs. Fig. 3(a) shows a SU8686 pancreatic ductal adenocarcinoma cell (acquired from ATCC, CRL-1837) labeled with Tractin-CyOPF1 as imaged with our OPM and after deconvolving the data. Importantly, the cross-sectional view revealed excellent optical sectioning and enough resolving power to distinguish surface ruffles. In fig. 3(b-c), renderings of the segmented SU8686 cell from two viewpoints are shown, with the color indicating surface intensity. Fig. 3(d) shows a spheroid made of colorectal adenocarcinoma cancer cells DLD1 labeled with tractin-GFP as imaged with our microscope and after deconvolving the data (cell culture description in the Section 6 of Supplement 1). Fig. 3(e-f) shows renderings of the segmented surface from two viewpoints, with the color indicating surface intensity. The 3D segmentation and surface rendering are discussed in detail in Section 7 of Supplement 1. Time-lapse imaging of the spheroid is shown in Supplementary Movie 1, which reveals rapid protrusion dynamics that are resolved spatially and temporally.

**Fig. 3.**
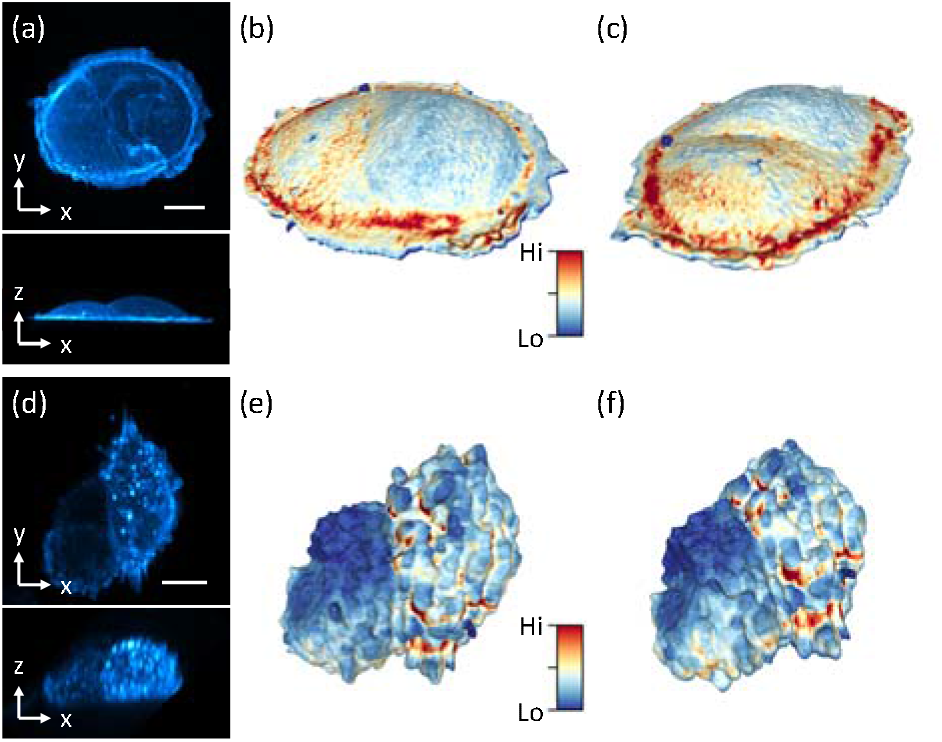
Imaging of single cells and spheroids. (a) Maximum intensity projection (MIP) of SU8686 cell in xy and xz. (b-c) 3D rendering of cell surface from two different views. Color encodes surface intensity (d) MIPs of Colon cancer cell spheroid in xy and xz. (e-f) 3D rendering of the surface from two different views. Scale bars: (a) 20 µm, (d) 10 µm.

To demonstrate the potential of optical tiling OPM for high resolution intravital microscopy, we performed longitudinal imaging of the vascular development in the zebrafish tail. For this, a 1.5 days old zebrafish labeled with the vasculature marker Tg(kdrl:Hsa.HRAS-mCherry)[16] was imaged over a time span of 16 hours (the design of water reservoir for long-term imaging is shown in Fig. S11 in Supplement 1). To visualize the majority of the tail vasculature at each timepoint, three optically tiled stacks were acquired and computationally fused together using BigStitcher [17] (Fig. 4a, Supplementary Movie 2). The high resolving power of our system enabled us to visualize the intricate process of tip cell migration of the nascent intersegmental vessel (ISV), the active extension of numerous endothelial filopodia that dynamically probe the environment, and initiation of anastomosis (vessel fusion) between neighboring sprouts, resulting in a new vessel, the dorsal longitudinal anastomotic vessel (DLAV) (Insets in Fig. 4b).

**Fig. 4.**
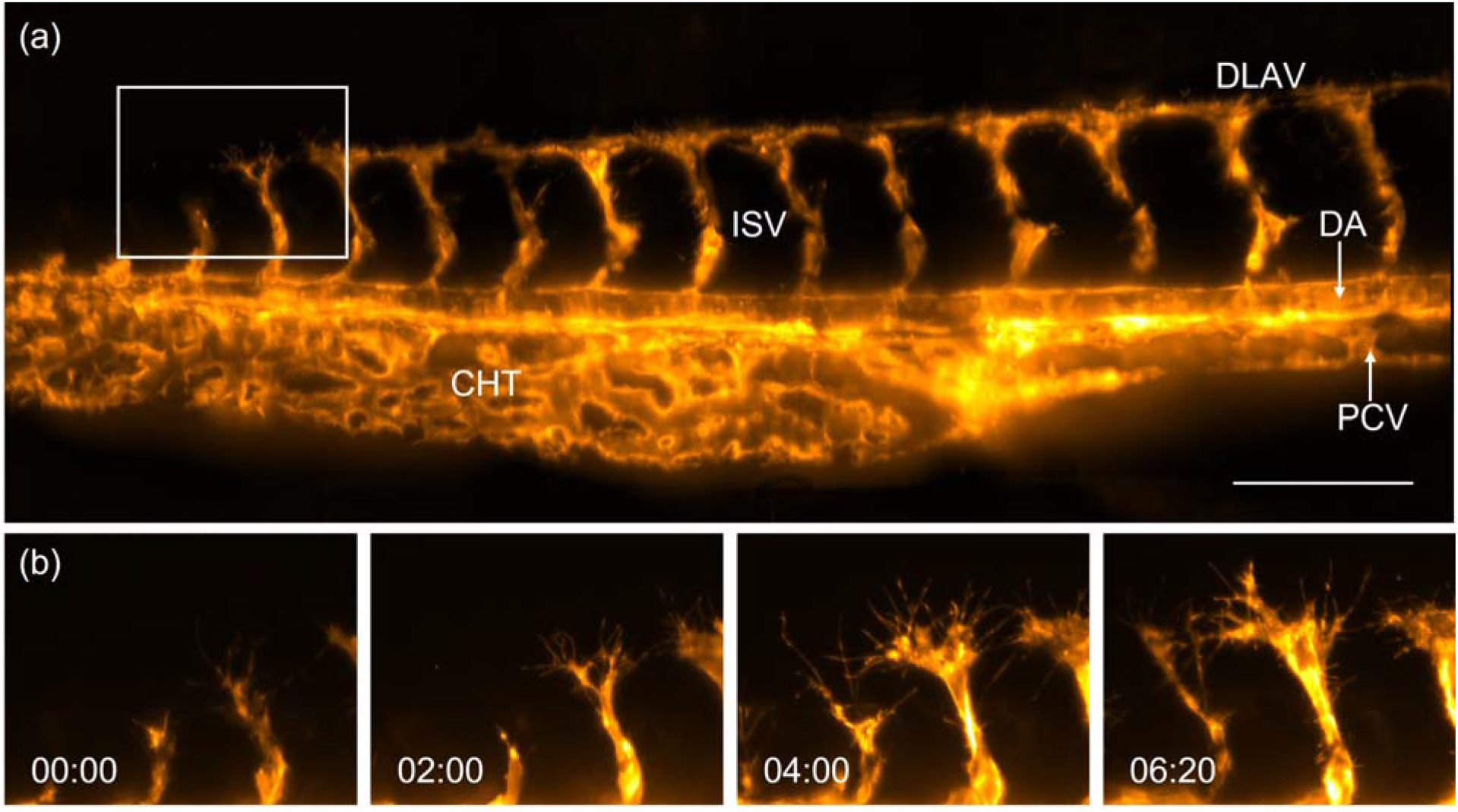
Imaging of zebrafish vasculature with optical tiling OPM. (a) Tail of a 1-2 d old Zebrafish labeled with Tg(kdrl:Hsa.HRAS-mCherry), as imaged by optical tiling OPM. A maximum intensity projection of three volumes that have been computationally stitched together is shown. DLAV, Dorsal longitudinal anastomotic vessel; DA, Dorsal aorta; PCV, Posterior cardinal vein; ISV, Intersegmental blood vessel; CHT, caudal hematopoietic tissue. (b) Enlarged views of the boxed region in (a) show the development of intersegmental vessels in the zebrafish tail. Scale bar: 100 µm. Timestamp: hh:mm.

Lastly, we demonstrate that our optical tiling technique can be used to extend the FOV in applications that require acquisition speeds that lie outside the timescales accessible by conventional 3D stack acquisitions. One promising approach to do so is the recently introduced real-time multi-angle projection method [14]. Fig. 5(a) shows the schematic principle of the projection mechanism: The light-sheet is rapidly swept through the sample (completing at least one sweep per camera exposure). Synchronized to this sweep, the shear galvo is translating the instantaneous images across the image sensor. This results in an optical implementation of the shear-warp transform, yielding projections of the sample under variable viewing angles [Fig. 5(a), bottom]. As an example, six different projections of vasculature in the zebrafish tail, labeled with the vascular marker Tg(kdrl:EGFP) [18], are shown in fig. 5(b). By rapidly tiling three projections, we imaged the beating heart of a zebrafish, labeled with the vascular marker Tg(kdrl:EGFP) and the red blood cell marker Tg(gata1a:dsRed) [19] in a casper background [20]. We achieved an overall acquisition rate of 13 Hz using 3 optical tiles, projected over a depth of 100 microns [Fig. 5(c), with time coded by color, and supplementary Movie 3].

**Fig. 5.**
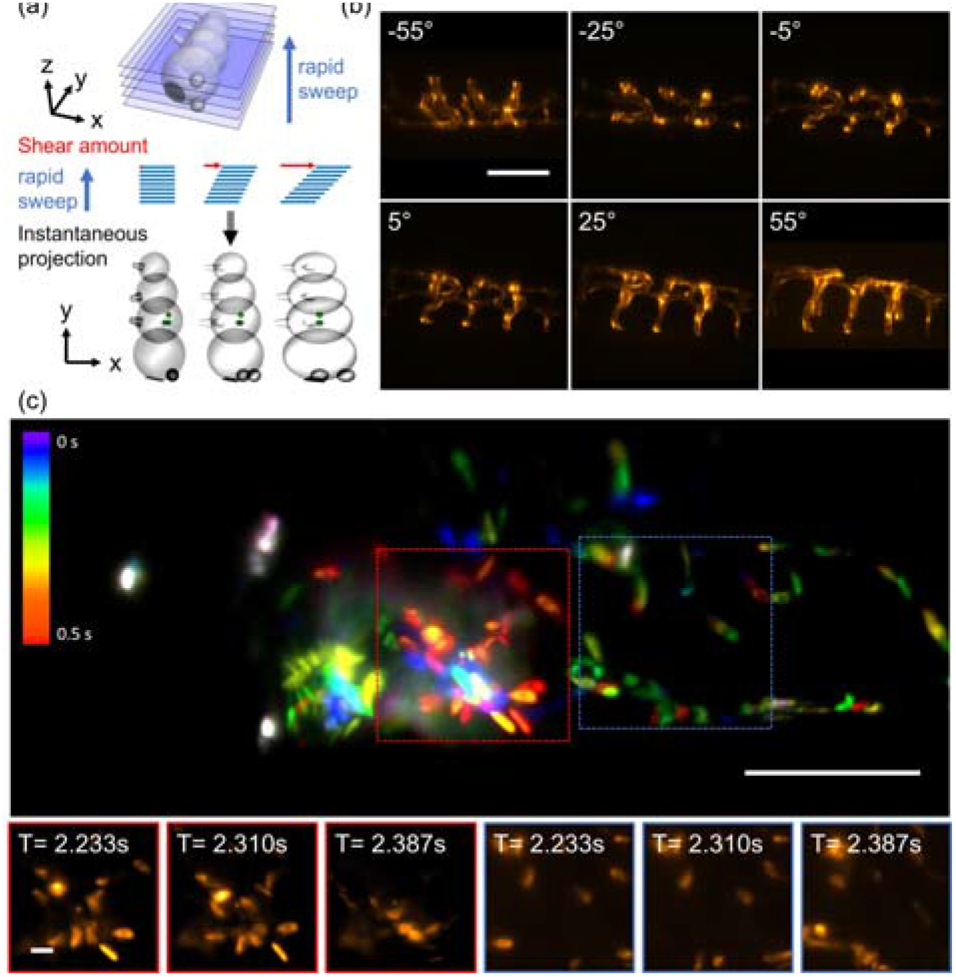
Extending the field of view in real time projection imaging. (a) Schematic representation of the multi-angle projection method: the light-sheet is rapidly swept through the sample, and the instantaneous images are translated at the same rate on the camera sensor. Depending on the amount of lateral translation, projections under different viewing angle are obtained. (b) Projections of vasculature in the zebrafish tail under different viewing angles. (c) tiled projection imaging of fluorescently labeled blood cells in a zebrafish heart. Insets show a montage of blood cells at different timepoints. Scale bars: (b-c) 100 µm and insets 10 µm.

In summary, we have introduced a method to extend the FOV in OPM via optical tiling. This overcomes a common bottleneck in OPM systems, as the FOV of the tertiary imaging system is demagnified into the sample space. Here, we exploit the fact that the primary lens in an OPM system can provide a large FOV and high spatial resolution, which we in turn made accessible with a custom dual-axis scanning unit. Importantly, this scanning unit does not require any additional relay lenses; it fits in the same space where a single galvo would be located in a conventional OPM system. Our system covers a field of view of 800 × 500 microns, a 2.5-fold improvement over a conventional OPM scanning approach, at subcellular resolution while providing Nyquist sampling.

In contrast to mechanical tiling, our optical implementation is much more rapid (on the order of 1-2 milliseconds) and does not induce sample vibrations. Applications include observing large samples without mechanical perturbation and at high speed, and potentially follow a region of interest dynamically. As demonstrated by our imaging experiments, the presented OPM system provides enough spatial resolution to resolve fine subcellular features such as clathrin coated vesicles, cell ruffles, blebbing and sprouting of endothelial cells. It also features the temporal resolution to follow rapid dynamics, such as the blood flow through a beating zebrafish heart. Notably, mechanical tiling would not have been compatible with the framerates used in the real time projection application. As such, we think that optical tiling OPM, with its ability to provide high spatiotemporal resolution over extended FOVs, will find applications from single cell studies over organoids to model organisms.

## Data availability

Data may be obtained from the authors upon reasonable request.

## Supplemental document

See Supplement 1 for supporting content.

