## Supplemental Document for "Increasing the Field-of-View in Oblique Plane Microscopy via optical tiling"

This document provides the details of the dual-axis scan unit, the single-galvo shear unit, the illumination module, the performance of the ASLM mode, the 3D segmentation and meshing of volumetric data and sample preparation.

1. Dual-axis scan unit

We used Zemax and geometrical optics to design, analyze and present the working principle of the dual-axis scan unit in this section. Fig. S1(a-c) shows the 3D layout of the Zemax simulation model with 9 configurations (shown in different colors), which correspond to the output optical scanning angle of 0° and ±5° in each direction and could support the FOV of 12.2 x 12.2 mms after the scan lens for point scanning. Fig. S1(d) shows a rendering of the CAD model for the dual axis scan unit.

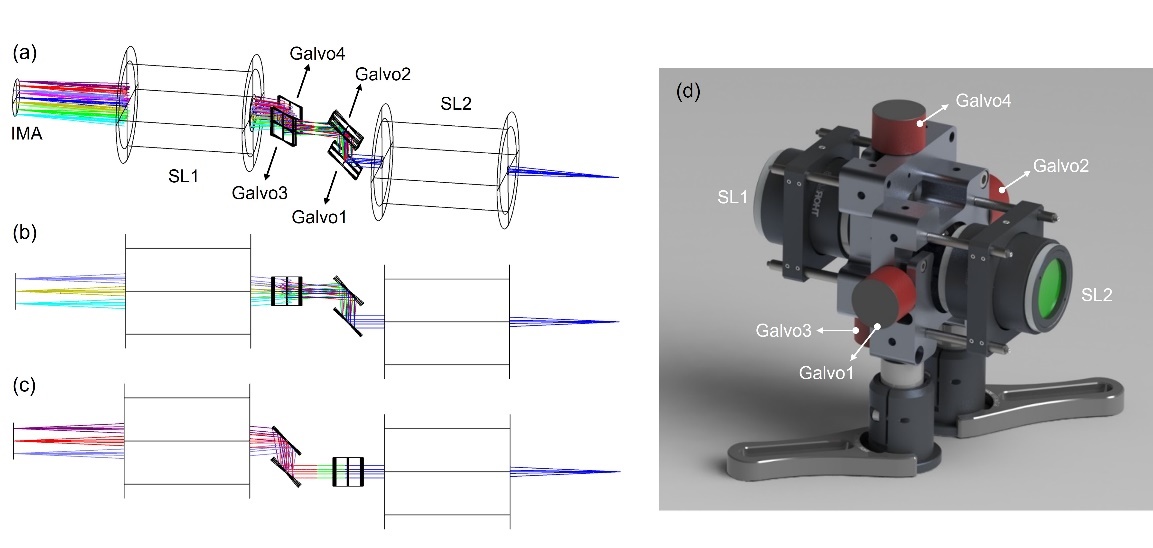

Fig. S1. (a-c) 3D Layout of the dual axis scan unit and the adjacent scan lenses. SL1-2, Scan Lenses; IMA, image plane. Light is coming from the right. Tilted view (a), Side view (b), Top view(c). (d) CAD model of the mechanical assembly including the scan unit and the scan lenses.

Fig. S2(a) shows the spot diagram of 9 configurations at image plane [IMA, in Fig S1(a)]. The black circle indicates the airy radius of around 8.37 µm and the RMS spot size in all configurations are well under the Airy radius. To compare the performance with a conventional scan strategy, we replaced the dual-axis scan unit with two Galvos overlapped at the focal plane of the scan lens in Zemax [shown in Fig S2(b)], which cannot exist in reality. The Huygens PSFs of the lower left configuration with the dual axis scan unit and ideal two-galvo scan unit are shown in Fig.S2(c) and (d). The corresponding Strehl ratios are 0.766 and 0.765, which demonstrates that the distortion shown in the spot diagrams is from the current chosen scan lenses, not from the design of the dual axis scan unit.

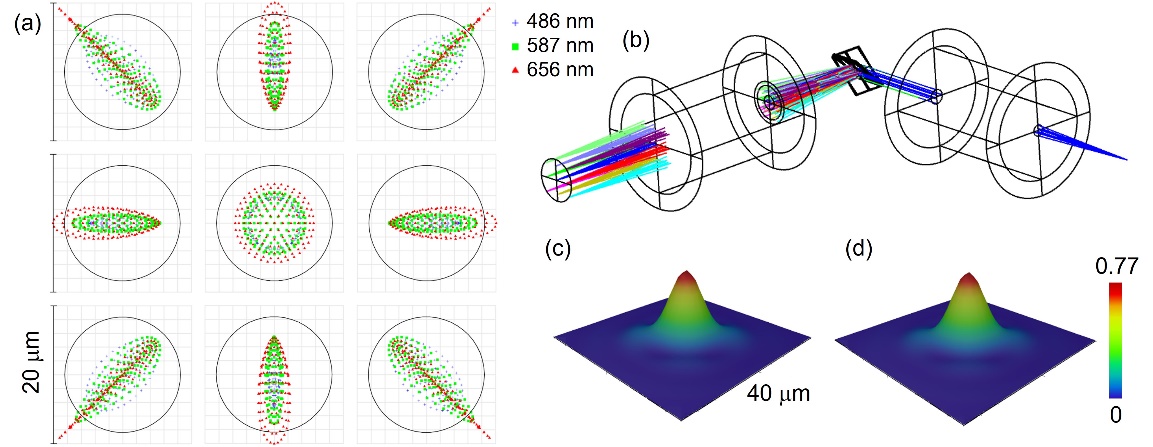

Fig. S2. (a)Configuration matrix spot diagram of the Zemax model for the dual axis scanner. Here the chief ray is used as reference. Legend items refer to wavelength. (b) 3D Layout of the ideal two-galvo scanner with the same scan lenses. (c) Huygens PSF of the lower left configuration with the dual axis scan unit. (d) Huygens PSF with the ideal two-galvo scanner.

The optical scanning angle of our dual axis scan unit with the Thorlabs GVS211 galvanometric mirrors is limited to ±5°. The Zemax simulation illustrates ray clipping of the galvo at the extreme scan position, which causes the degradation of resolution in x axis (Fig. 2(a)). As shown in Fig. S3(a) and Fig. S3(b), the three beams in different colors indicate three points (center and both ends) illuminated by the light sheet at the optical scanning angle of ±5° [see also green volumes in Fig.1(g)]. The green light in Fig. S3(c) and the red light in Fig. S3(d) show ray clipping at both ends. Of note, the beam clipping depends on the beam size in the simulation, thus on the size of Back Focal Plane (BFP) of the objective. It could be alleviated by using the objectives with smaller BFP size or using larger Galvo mirrors as Galvo 2 and 3 in the scan unit.

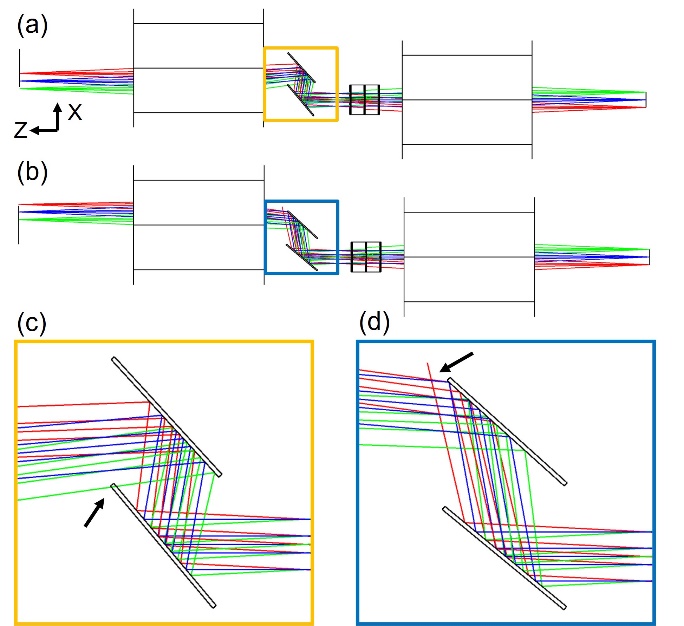

Fig. S3. (a) 3D Layout of the dual axis scan unit and the adjacent scan lenses in the light-sheet condition with the optical tiling angle of 5°. (b) 3D Layout with the optical tiling angle of -5°. (c, d) Zoom-in of the box in (a) and (b), respectively. Arrows indicate the ray clipping.

The Zemax simulation shows the feasibility of the dual-axis scan unit. We further use geometrical optics to demonstrate the relation between two Galvos and the common point in each Galvo pair. Fig. S4(a) and (b) show the working principle of Pair 1 and Pair 2, which correspond to Fig.1(d) and (e).

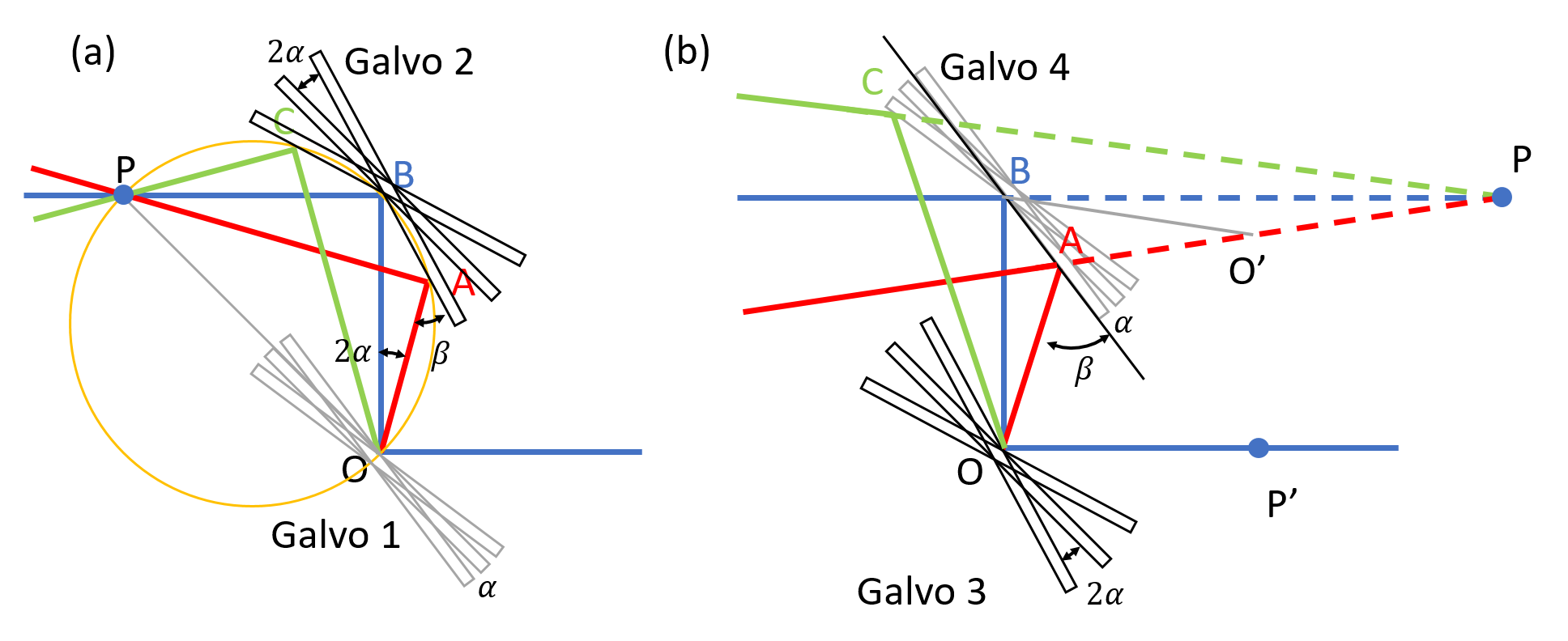

Fig. S4. Working principle of dual-axis scanner. (a) Galvo pair 1. (b) Galvo pair 2. P’ is the common point P in (a).

For Pair 1[shown in Fig.S4(a)], we first demonstrate that if the scanning angle of Galvo 2 is twice the scanning angle of Galvo 1, all laser beams will pass through a common point P. This can be proved by showing that points A, B, O, P are on one circle (P is the intersection point of laser beams with the scan angle of 0 and $2\alpha$). Then, we can show the distance between the common point P and Galvo 2 equals to the distance between the two Galvos, which means OB = BP.

If we rotate Galvo 1 for $\alpha$ to get the red beam, the optical scanning angle $\angle AOB$ will be $2\alpha$. If we then rotate Galvo 2 for $2\alpha$, we have

$\angle ABO=45^{\circ}-2\alpha$.

$∵\beta=\angle ABO+\angle AOB=45^{\circ},$ $\angle BAP=\beta$,

$\therefore\angle OAP=90^{\circ}$.

We already know $\angle OBP=90^{\circ}$, then $\angle OAP$=$\angle OBP$. According to the inscribed angle theorem, we can demonstrate that A, B, O, P are on one circle.

Because points A, B, O, P are on one circle, we have

$\angle APO=\angle ABO=45^{\circ}-2\alpha$.

Then $\angle AOP=90^{\circ}-\angle APO=45^{\circ}+2\alpha$, $\angle BOP=$ $45^{\circ}$. So $\Delta OBP$ is an isosceles right triangle and OB = BP.

Similarly, we can prove P is also the intersection point of laser beams with the scan angle of 0 and -$2\alpha$ (blue and green).

Therefore, all laser beams after Pair 1 will pass through the point P, and OB = BP.

Next, we can prove for Pair 2, when the scanning angle of Galvo 3 is set to twice the scanning angle of Galvo 4, a common virtual scan point P can be formed, as shown in Fig. S4(b).

If we rotate Galvo 3 for $2\alpha$ to get the red beam, the optical scanning angle $\angle AOB$ will be $4\alpha$. Then if rotate Galvo 4 for $\alpha$, we have

$\angle ABO=45^{\circ}-\alpha$,

$\angle BAO=180^{\circ}-\angle ABO-\angle AOB=135^{\circ}-3\alpha$.

$∵\angle ABP=45^{\circ}+\alpha$*,* $\angle BAP$*=* $\angle BAO=135^{\circ}-3\alpha$,

$\therefore\angle APB=2\alpha$.

We can add point $O^{'}$, which makes ${AO}^{'}=AO$ and we can easily prove that $\Delta ABO'$ is congruent to $\Delta ABO$. Then we have $\angle AO^{'}B=\angle AOB=4\alpha$, $\angle O^{'}BP=$ $90^{\circ}-2\left( 45^{\circ}-\alpha\right)=2\alpha$, and consequently $\angle{BPO}^{'}=2\alpha$. Thus, $\Delta BO'P$ is an equilateral triangle, $O^{'}B=O^{'}P$, and

$$PB=2\cdot OB\cdot\cos\left( 2\alpha\right).$$

According to the equation shown above, the distance between the virtual point P and Galvo 4 is irrelevant to the sign of $\alpha$. Therefore, if Galvo 3 and 4 are rotated by $-2\alpha$ and $-\alpha$, respectively, shown as the green line in Fig S4(b), the above equation stays valid. This conclusion can also be proved by similar geometric analysis. Besides, we can also notice that when α is a small angle, the distance between the virtual point P and Galvo 4 is two times the distance between the two Galvos, $PB\approx2\cdot OB$. But at a larger angle, PB is dependent to α, which means there is a little deviation of the position of the virtual point P. We derive the deviation as

$\sigma PB=2\cdot OB\cdot[1-\cos\left( 2\alpha\right)]$.

In our case, $\sigma PB=2\times15 mm\times\left[ 1-\cos\left( 2\times5^{\circ} \right) \right]=0.456 mm$. This will cause the light-sheet to be slightly tilted away from the design oblique plane after the primary objective by $1.65^{\circ}$ at the edge, which is neglectable in our application.

[Derivation of the light-sheet tilt angle: $\mathrm{atan}\left( \sigma PB\times\frac{\tan\left( 2\times5^{\circ} \right)}{f_{SL}} \right)\times M=1.65^{\circ}$, where $f_{SL}$and $M$ are the focal length of scan lens and the magnification of the primary objective, $f_{SL}=70 mm$, $M=25$.]

2. Single-Galvo shear unit

Adding a pair of Galvo mirrors in front of the camera to shear the image has been demonstrated in [1] and is schematically shown in Fig. S5(a). Here we adopted a simpler implementation with a single-Galvo scanner as shown in Fig. S5(b) and demonstrate its virtually identical performance.

For both shear units, the optical path length will change for different shearing amounts, which in principle adds a small amount of defocus to the image. In addition, the single-Galvo shear unit adds a small amount of tilt of the image as it is being scanned, which compresses the images in the shear direction. These effects however turn out to be neglectable based on our analysis.

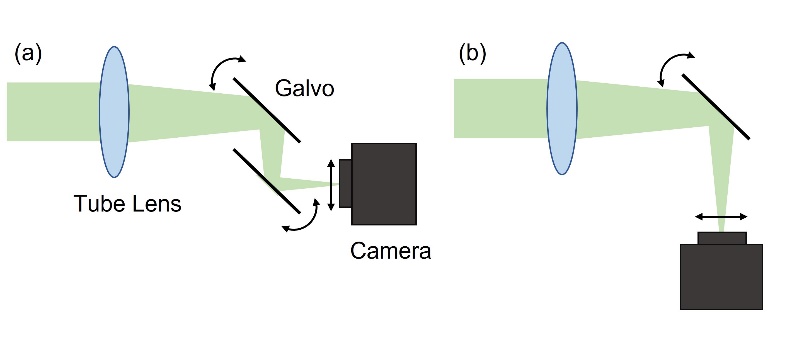

Fig. S5. Schematic of the shear unit with (a) dual Galvo and (b) single Galvo.

To calculate the variation of the optical path length at the surface of the camera chip, we drew the optical axis of convergent fluorescence light without shearing (blue lines, shown in Fig. S6) and with the shear angle $\alpha$ (red lines). The shear angle will shift the optical axis by x at the surface of the camera chip (dashed lines). The length difference of the red lines and blue lines is the light path difference.

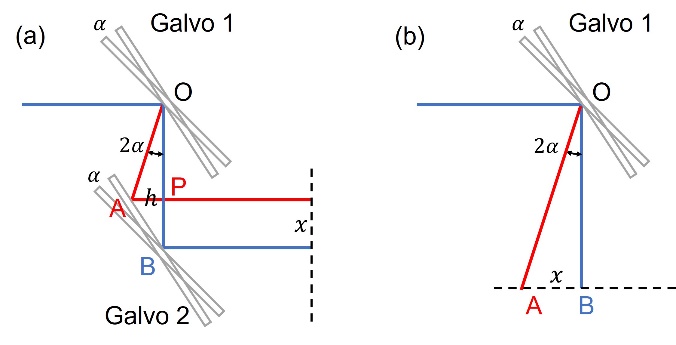

Fig. S6. Working principle of dual-Galvo shear unit (a) and single-Galvo shear unit (b).

For dual-Galvo shear unit as shown in Fig. S6(a), the light path difference is OA+AP-OB. When the Galvo rotated by $\alpha$, the optical scanning angle is $2\alpha$, then we have $\angle AOB=2\alpha$, $\angle ABO=45^{\circ}-\alpha$. Defining $OB=d$, $AP=h$, we have

$$h=OP\cdot\tan\left( 2\alpha\right)=PB\cdot\tan\left( 45^{\circ}-\alpha\right),$$

$$OP+PB=OB,$$

$$\therefore OP+PB=\left( \frac{\tan\left( 45^{\circ}-\alpha\right)}{\tan\left( 2\alpha\right)}+1 \right)x$$

$$=\frac{x}{\sin\left( 2\alpha\right)}=d.$$

Therefore, the light path difference for dual-Galvo unit is

$$OA+AP-OB=\frac{h}{\sin\left( 2\alpha\right)}+h-d=x\tan\left( 45^{\circ}-\alpha\right)\left( \frac{1}{\sin\left( 2\alpha\right)}+1 \right)-d=-2\sin^{2}\left( \alpha\right)d=-2d\sin^{2}\left( \frac{1}{2}\mathrm{asin} \left( \frac{x}{d} \right) \right).$$

For single-Galvo shear unit, the light path difference is

$$OA-OB=\sqrt{x^{2}+d^{2}}-d.$$

Therefore, we can get the focal plane deviation with the camera chip as abscissa under different conditions, shown in Fig. S7(a). For the dimensions chosen in our setup, *i.e.* the galvo is placed 110 mm in front of the camera in the single-Galvo shear unit (OB in Fig. S7(b)), and the distance between two galvos in the dual-Galvo shear unit is 25.4 mm (OB in Fig. S7(a)), the maximum focal plane deviation is around 0.2 mm for single-Galvo shear unit and -0.88 mm for dual-Galvo shear unit. For -0.88 mm deviation, the corresponding focal shift in sample space is 0.23 µm, which is still below the depth of focus of the OPM imaging system. Nevertheless, if we extend the distance between the galvos in the dual-Galvo shear unit, we can see the deviation is close to the single-Galvo shear unit (orange line in Fig. S7(a)). Given the large distance from the single-Galvo shear unit to the camera chip, the tilted optical axis will lead to a 99.8% compression of the images at the extremes of the scan range. The compression is the ratio of the projection of the tilted image to the un-tilted image, which is a simple cosine relation and is cos(3.46°) in this case. The focal plane with equal light path length (blue lines) at different shear angle is shown in Fig. S7(b) and (c).

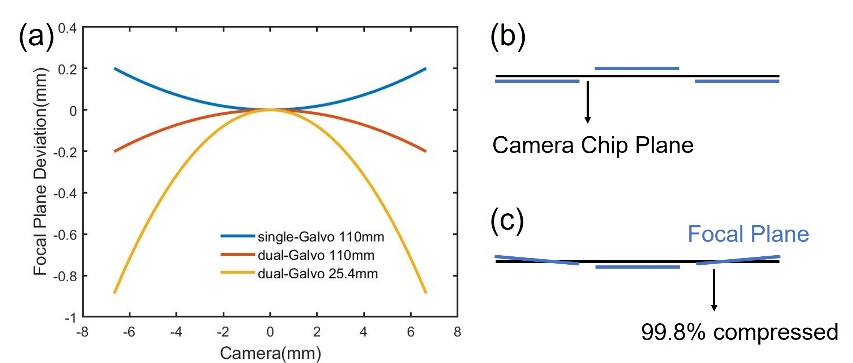

Fig. S7. (a) Focal plane deviation at camera chip. Schematic of focal plane deviation for (b) dual-Galvo unit, and (c) single-Galvo unit.

3. The illumination module

A schematic drawing of the illumination module is shown in Fig S8(a). The output laser from a fiber coupled solid state laser module (OBIS Galaxy with laser modules LS 561nm-80mW, LX 488nm-100mW and LX 640nm-75mW, Coherent Inc) was first collimated by a fiber collimator (CFC11A-A, Thorlabs), then expanded in one dimension by a Powell lens and an achromatic lens L1 (2x AC254-030-A, Thorlabs) (shown in “side-view”). A telescope of achromatic doublets L2 (AC254-060-A, Thorlabs) and L3 (AC254-050-A, Thorlabs) images the light-sheet on a resonant galvo (CRS 4KHz, Cambridge technologies), which is conjugate to the image plane of the OPM system. After the resonant scanner, an achromatic doublet L4 (AC254-040-A, Thorlabs) forms a 4F system with the tube lens TL2 [see also Fig.1(a)] to conjugate the resonant scanner to the image plane of the OPM system. As shown in the top-view, an ETL(Electrically tunable lens, EL-16-40-TC, Optotune) is placed on the Fourier plane between L2 and L3 to refocus the waist of the light-sheet and the beam condition will be maintained in the other dimension. An adjustable slit is used to control the divergence of the light-sheet. The mirror after L4 is conjugate to the pupil plane of the primary objective. As such, changing the mirror tilt angle translates the light-sheet in the sample plane. We used a Gimbal Mirror Mount (U100-G2K, Newport) to ensure only the angle of the beam is tuned and no shift is introduced. We further replaced the manual adjustor of the mount with a piezo actuator (PIA25, Thorlabs) to allow motorized fine-tuning capability. The illumination unit sits on a large translation stage (#66-455, Edmund optics). When translating the whole illumination unit, the light-sheet incident angle can be varied in fine steps. Fig. S8(b) shows photo of the optical setup. The system is built to allow the ETL lying horizontally.

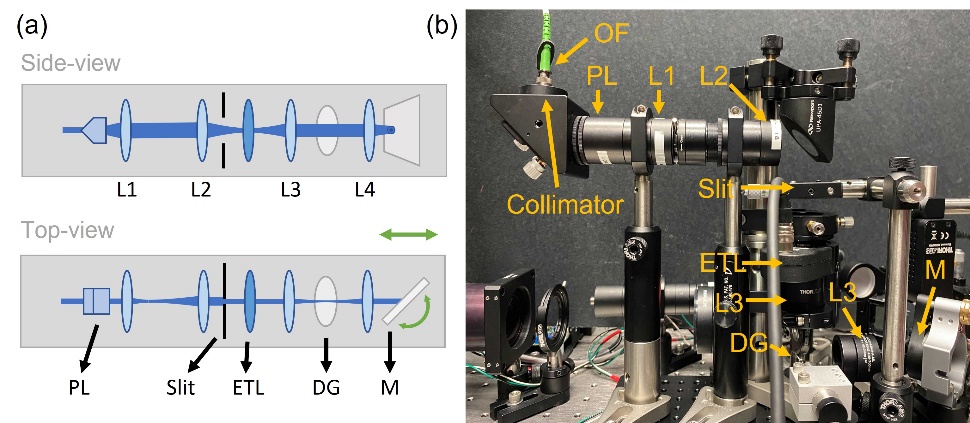

Fig. S8. (a)Schematic of the illumination unit in side-view and top-view. (b)Photo of the illumination unit. L1-4, Lenses; PL, Powell Lens; ETL, Electrically Tunable Lens; DG, Dither Galvo; M, Mirror; OF, Output fiber.

4. Axially Swept Light-Sheet Microscopy imaging mode

In standard light-sheet microscopy, a long and thick light-sheet is needed to cover a large sample, which results in a worse axial resolution. Axially Swept Light-Sheet Microscopy (ASLM) overcomes this challenge by axially sweeping a short, thin light-sheet and synchronizing the sweep of the light-sheet with the rolling shutter of the camera. With an ETL in our illumination unit, we can realize an ALSM imaging mode in OPM. We first used fluorescent nanospheres to measure the resolution [Fig. S9(a)]. For the thin light-sheet, the axial resolution (measured by the FWHM of the nanospheres in z) was around 1 µm at the waist of the light-sheet and increased to around 2 µm at the edge, while for the ASLM, an axial resolution of 1.01±0.14 µm was maintained throughout the volume. The axial resolution gain is modest, as the available NA for the light sheet is more limited than in a conventional ASLM system. However, the ASLM mode can help to homogenize axial resolution in an OPM system.

We further tested the performance of the ASLM in live sample by imaging the AKP (APC^-/^; KRAS^G12D/+^; p53^-/-^; TdTomato^+^) organoids [2] in both modes [Fig. S9(b)]. The organoid cells are labelled by TdTomato and is visualized in magenta and β-catenin is labelled by 488 AlexaFluor and shown in green.

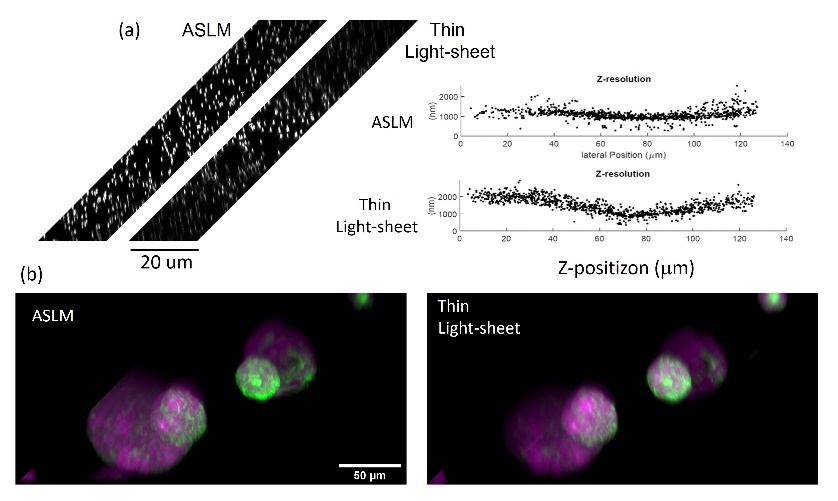

Fig. S9. (a) Fluorescent nanospheres (shown in y-z view) and their Full Width Half Maximum measurements in ALSM mode and thin light-sheet mode. (b)AKP Organoids imaged in ASLM mode and using a thin light-sheet.

5. Deconvolution and resolution analysis

To evaluate the resolution in a live cell imaging context, we preformed image decorrelation analysis [3]on ARPE cells labeled with EGFP for AP2 before [Fig. S10(a)] and after [Fig. S10(b)] deconvolution. Before the deconvolution, we up sampled the image by a factor of 2 through zero padding at the Fourier Transform image. Using image decorrelation analysis on the whole image, the lateral resolution is 408 nm [Fig. S10(c)] in the raw data, and 294 nm [Fig. S10(d)] after deconvolution.

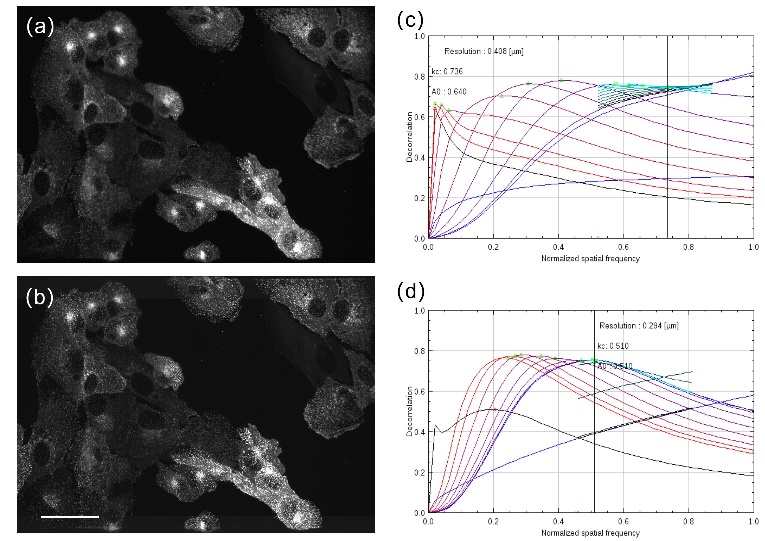

Fig. S10. (a) Maximum projection of parental human retinal pigmented epithelium (ARPE-19) cells with EGFP-labeled AP2. (b)After up-sampling and deconvolution. (c)Decorrelation analysis of a plane of (a). (d) Decorrelation analysis of the same plane in (b).

6. Colon cancer cell spheroid culture

The colorectal adenocarcinoma cancer cells (DLD1, colon cancer cell line) in spheroid form were made as follows:

Day 1:

1. Warm 1.5% Difco Noble Agar (in PBS) in microwave until it is completely dissolved.
2. For a 96 well plate, add 50 µL of 1.5% Difco Noble Agar to each well using a 12-channel pipette and a reservoir. (Make sure there are no bubbles)
3. Allow agar to set at rt.
4. Bring up cells in media with 10% FBS. Count cells. (Can first try media used to culture cells, changing % FBS can help spheroids form)
5. Plate 5,000 cells/well in 200 µL of media. Incubate at 37º C until spheroids form (3-5 days).

Day 4-7:

Pipette up gently to release spheroid from agar. Transfer media with spheroids to a labeled 15 mL conical tube. Let sit at rt. until spheroids fall to the bottom of the tube.

Discard media carefully and seed organoids on a layer of matrigel or 1% collagen.

7. Segmentation and meshing of 3D imaging volume

To segment and extract the surface mesh, the raw volumetric image intensity is first min-max normalized to [0-1] and contrast stretched between 2% and 99.8% percentile to enhance image features. Contrast limited adaptive histogram equalization (CLAHE) is applied to locally enhance contrast with a kernel size 1/8^th^ of the original volumetric image shape and a clipping intensity limit of 0.01. The result is normalized by contrast stretching between 2% and 99.8% percentile. Maximum likelihood blind deconvolution was applied for 30 iterations to learn a PSF given an initially synthesized PSF. The learnt PSF was then used to deconvolve the image using Wiener-Hunt deconvolution, balance=1, and Laplacian regularization. The deconvolved image is clipped to [0-1], contrast stretched between 2% and 99.8% percentile and standard normalized. The image is clipped between 0 and 4 standard deviations and min-max scaled to [0-1] to obtain the final postprocessed image intensities for segmentation. From this image we obtain the final binary segmentation volume image by deriving and combining three different segmentations. The first segmentation is obtained from Otsu threshold and captures the majority of the cellular detail. The second segmentation is optimized to capture a smooth inner core shape of the cell/organoid. It is obtained by significant oversmoothing with a Gaussian filter of the postprocessed image, and thresholding with mean+s.d. with morphological hole filling. The third segmentation is optimized for the detection of thin, high-frequency protrusions. It is obtained by computing the difference of Gaussian (DoG) image which enhances image edges given by postprocessed image – gaussian filter (postprocessed_image, sigma=3), then computing and thresholding the normalized DoG image ((DoG-mean(DoG))/4std(DoG) >= 1). The final binary image is smoothed with a Gaussian filter of sigma=1 and meshed at a contour level of 0.5 using marching cubes. The final mesh is obtained after remeshing the marching cubes mesh to obtain a uniform triangular mesh with approximated centroidal voronoi diagram (ACVD) after collapsing small edges. Fluorescence intensities were then mapped onto the mesh by trilinear interpolation at vertex coordinates. The RdYlBu lookup table was used to assign colors to intensity values using 1% and 99% intensity values as the lower and upper clipping limits. Colored meshes were saved as .obj files and rendered using Meshlab. For the timelapse movie, volumetric images were rigid registered temporally using frame 0 as the common reference image and intensities corrected for bleaching with histogram matching to the frame 0. The segmentation described above was applied to obtain meshes for each timepoint.

8. Water reservoir for long-term imaging

The used primary objective has a 2 mm working distance. Due to the inverted geometry of our setup, this requires a long water meniscus, which is prone to evaporation when performing long-term imaging. We used a water reservoir below the coverslip to minimize water evaporation and provide a larger water basin, as shown in Fig. S11.

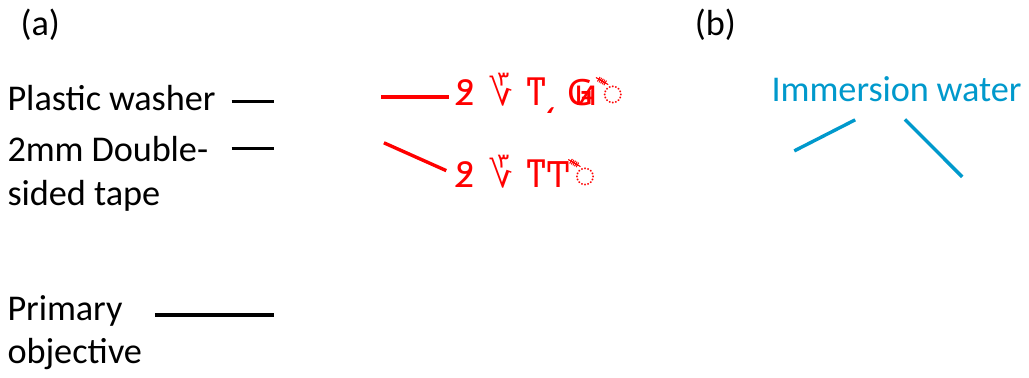

Fig. S11. Water reservoir for long-term imaging. (a) The water reservoir is assembled by adhering a 2 mm thick double-sided tape (Amazon) and a plastic washer (Chemical-Resistant PTFE Plastic Washer, 94115K005, McMaster-Carr) on the primary objective. The double-sided tape is cute to a ~30 mm square with a ~ 10 mm hole. The plastic washer has an inner diameter of 0.532 inches. (b) A large amount of immersion water (compared to a conventional water meniscus) can be added. Although it still slowly vaporizes, water levels stay sufficiently high for imaging for at least 24 hours.
